# piRNA clusters need a minimum size to control transposable element invasions

**DOI:** 10.1101/838292

**Authors:** Robert Kofler

## Abstract

piRNA clusters are thought to repress transposable element activity in mammals and invertebrates. Here we show that a simple population genetics model reveals a constraint on the size of piRNA clusters: the total size of the piRNA clusters of an organism ought to exceed 0.2% of a genome. Larger piRNA clusters accounting for up to 3% of the genome may be necessary when populations are small, transposition rates are high and TE insertions recessive. If piRNA clusters are too small the load of deleterious TE insertions accumulating during a TE invasion may drive populations extinct before an effective piRNA based defence against the TE can be established. Our finding is solely based on three well supported assumptions: i) TEs multiply withing genomes, ii) TEs are mostly deleterious and iii) piRNA clusters act as transposons traps, where a single insertion in a cluster silences all TE copies in trans. Interestingly, piRNA clusters of some species meet our minimum size requirements while clusters of other species don’t. Species with small piRNA clusters, such as humans and mice, may experience severe fitness reductions during invasions of novel TEs, possibly even threatening the persistence of some populations. This work also raises the important question of how piRNA clusters evolve. We propose that the size of piRNA clusters may be at an equilibrium between evolutionary forces that act to expand and contract piRNA clusters.

## Introduction

Transposable elements (TEs) are short stretches of DNA that selfishly propagate within genomes (Orgel and Crick, 1980; Doolittle and Sapienza, 1980). It is thought that the proliferation of TEs is mostly deleterious to hosts. Negative effect of TEs may arise by three mechanisms: i) ectopic recombination among TEs could lead to deleterious chromosomal rearrangements ii) TE insertions may have direct negative effects, for example by disrupting genes or regulatory regions and iii) the products of TEs, such as the *Transposase*, could generate deleterious effects (e.g. DNA breaks) (Nuzhdin, 1999; Montgomery et al., 1991). Despite this largely selfish activity some TE insertions may confer beneficial effects to hosts, such as resistance to insecticides (González et al., 2008; Casacuberta and González, 2013; Daborn et al., 2002). However, the distribution of fitness effects of TE insertions is an important open question (Arkhipova, 2018; Kofler, 2019). An unrestrained proliferation of TEs could drive host populations extinct (Kofler, 2019; Brookfield and Badge, 1997), hence the spread of TEs needs to be controlled. The proliferation of TEs may be controlled at the population level by negative selection against TE insertions and at the host level, for example by small RNAs that repress TE activity (Charlesworth and Charlesworth, 1983; Charlesworth and Langley, 1989; Brennecke et al., 2007; Blumenstiel, 2011; Kofler, 2019). In mammals and invertebrates the host defence systems relies on piRNAs, small RNAs with a size between 23-29nt (Brennecke et al., 2008; Gunawardane et al., 2007). These piRNAs bind to PIWI-clade proteins that direct repression of TEs at the transcriptional and post-transcriptional level (Gunawardane et al., 2007; Brennecke et al., 2007; Le Thomas et al., 2013; Sienski et al., 2012). piRNAs are derived from discrete genomic loci, the piRNA clusters, which may make up substantial portions of genomes. In Drosophila, for example, piRNA cluster account for 3.5% of the genome (Brennecke et al., 2007).

It is thought that the proliferation of an invading TE is stopped when one copy of the TE jumps into a piRNA cluster, which triggers production of piRNAs against the TE that silence all TE copies in trans (Bergman et al., 2006; Malone and Hannon, 2009; Zanni et al., 2013; Goriaux et al., 2014; Yamanaka et al., 2014; Ozata et al., 2018; Duc et al., 2019). This view is known as trap model since piRNA clusters act as genomic traps for active TEs. The trap model is currently widely supported by many different observations. For example artificial sequences inserted into piRNA clusters yield piRNAs complimentary to the inserted sequence and a single insertion in a piRNA cluster is sufficient to silence a reporter construct in trans (Josse et al., 2007; Muerdter et al., 2012). Furthermore a study directly observing TE invasions noted a rapid emergence of piRNA cluster insertions and piRNAs complimentary to the invading TE (Kofler et al., 2018). Finally computer simulations confirmed that piRNA clusters may stop TE invasions (Kelleher et al., 2018; Kofler, 2019; Lu and Clark, 2009). Especially large clusters were able to control TE invasions under a wide range of different conditions (Kofler, 2019). It is however not clear if this observation holds for clusters of any size. Here we tested the hypothesis that piRNA clusters have a minimum size using computer simulations of TE invasions under the trap model. We found that piRNA clusters need to account for at least 0.2% of a genome to stop TE invasions over a wide range of parameters. Even larger piRNA clusters may be necessary when populations are small or TE insertions are recessive.

## Results

We performed forward simulations of TE invasions under the trap model to test the hypothesis that piRNA clusters need to have a minimum size. The trap model holds that the proliferation of an invading TE is stopped when a TE copy jumps into a piRNA cluster, which triggers the production of piRNAs that silence all TE copies in trans (fig. 1A). Accordingly we assumed that TEs multiply at a transposition rate *u* > 0 in individuals not having a cluster insertion and *u* = 0 in individuals having a cluster insertion. We simulated diploid organisms, with 5 chromosomes of size 10Mbp and a recombination rate of 4cM/Mb (fig. 1B). If not mentioned otherwise the population size was *N* = 1000. A piRNA cluster was simulated at the end of each chromosome (fig. 1B). We measured the size of piRNA clusters in percent of a genome, since we previously found that the relative size of piRNA clusters (in percent), and not the absolute size (in bp), determines invasion dynamics of TEs (Kofler, 2019). In our simulations piRNA clusters with a size of 1% correspond to one cluster of 100kb at the end of each chromosome (5 * 100*kb/*5 * 10*Mb*; fig. 1B). Initially we assumed that each TE reduces the fitness of the host (*w*) by a constant factor (*x*) such that *w* = *xn*, where *n* is the number of TE insertions per diploid (Charlesworth and Charlesworth, 1983). Furthermore we assumed that TE insertions in piRNA clusters incur no negative fitness effect (i.e. *x* = 0 for cluster insertions). To avoid the stochastic early stages of an invasion, where TEs may get lost due to genetic drift (Le Rouzic and Capy, 2005; Kofler, 2019), we launched each TE invasion by randomly distributing 1000 insertions in individuals of the starting population. These insertions have an initial population frequency of 1*/*2*N*.

**Figure 1:**
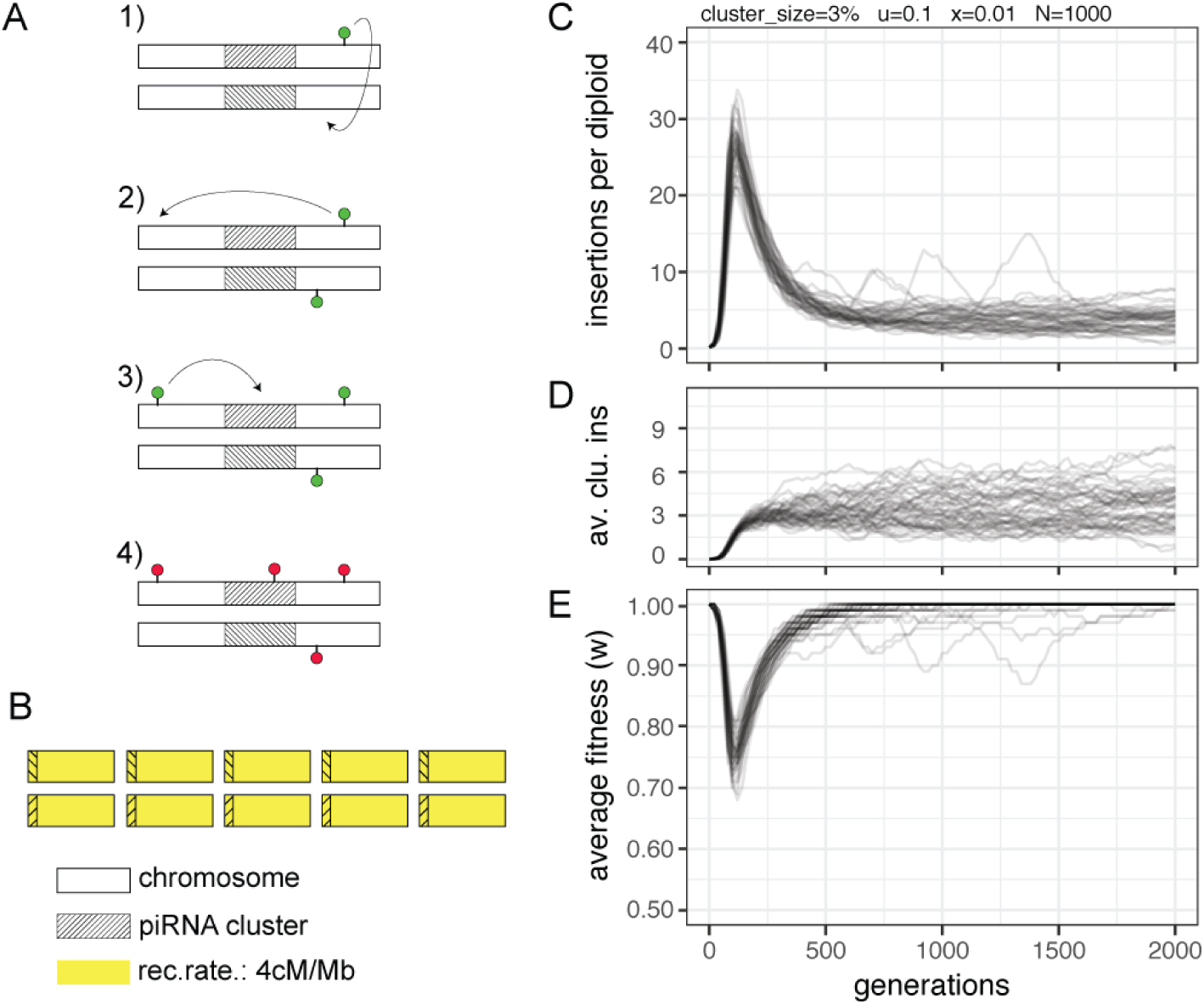
Dynamics of TE invasions with piRNA clusters and deleterious TE insertions. A) Under the trap model the proliferation of an active TE (green) is stopped when one copy jumps into a piRNA cluster (i.e. the trap; hatched area), which deactivates all TE copies in trans (red). B) We simulated five chromosomes with a size of 10Mbp for a diploid organism. A piRNA cluster was simulated at the end of each chromosome and a recombination rate of 4cM/Mb was used. C) Abundance of TE insertions during an invasion. Fifty replicates are shown. D) Number of cluster insertions per diploid during an invasion (av. clu. ins) E) Average fitness during an invasion.

Invasions of deleterious TEs under the trap model show a characteristic pattern (fig. 1C,D,E with *u* = 0.1, *x* = 0.01 and a cluster size of 3%). Initially the TE rapidly multiplies within a genome, which markedly reduces the fitness of the host (fig. 1C,E). Next individuals accumulate increasing numbers of cluster insertions, which slows down the spread of the TE (fig. 1C,D). Finally negative selection removes deleterious TE insertions, which restores fitness of the host nearly to initial levels (fig. 1C,E). In our example the TE invasion temporally reduced host fitness by about 20-30% (fig. 1E). In these simulations piRNA clusters accounted for 3% of the genome. The size of piRNA clusters is a major factor determining the amount of TEs accumulating during an invasion, where more TE insertions will accumulate when piRNA clusters are small (Kofler, 2019). We thus speculated that the fitness reduction during a TE invasion may be more dramatic for small piRNA clusters. Furthermore we surmised that populations may even go extinct when piRNA clusters are very small.

To test these hypothesis we simulated TE invasions with cluster sizes ranging from 0.001% to 10% (fig. 2A). Throughout this work we refer to the lowest fitness of a population during an invasion as the minimum fitness (fig. 2A). Furthermore we assumed that a population went extinct if its average fitness dropped below 0.1 (i.e. min. fitness < 0.1). With an average fitness of 0.1 about 16-23% of the individuals in a population have a fitness of zero (*w* = 0). These individuals will not contribute any offspring to the next generation. We introduced the extinction threshold of 0.1 to avoid unrealistic simulation conditions where, for example, some very few survivors with fitness just above zero could reconstitute the entire population of the next generation.

**Figure 2:**
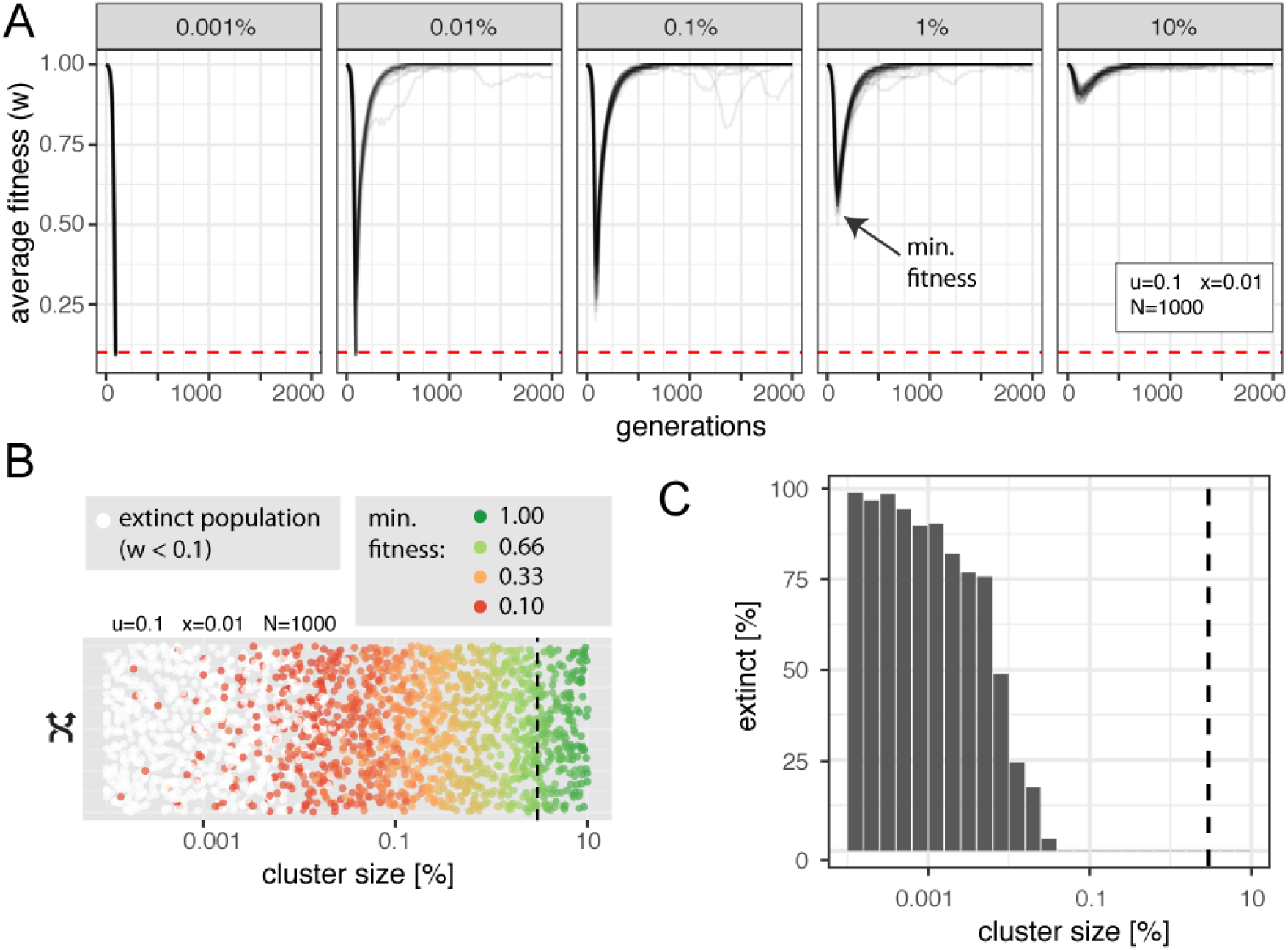
piRNA clusters need a minimum size to control TE invasions A) Average fitness of populations during TE invasions with different sizes of piRNA clusters (top panel). Fifty replicates are shown. We refer to the lowest fitness during an invasion as the “minimum fitness” (arrow). Red dashed lines indicate the extinction threshold (*w* < 0.1). With a cluster size of 0.001% all populations went extinct. B) Influence of the cluster size on extinction of populations. Each dot represents the result of single simulation with a randomly drawn cluster size. For non-extinct populations the minimum fitness is shown. The size of piRNA clusters in *D. melanogaster* is indicated as dashed black line (3%). C) Histogram showing the fraction of extinct populations for different cluster sizes. No extinct populations were found for clusters > 0.1%

As expected the minimum fitness was close to 1.0 for large clusters (fig. 2A; 10% cluster size). However, the minimum fitness dropped substantially with decreasing cluster sizes (fig. 2A). For clusters with a size 0.001% the minimum fitness dropped below 0.1 for all simulated populations, i.e. all populations went extinct. This demonstrates that transposon traps, such as piRNA clusters, need a minimum size to prevent extinction of populations. To identify the required size of piRNA clusters we performed 2000 simulations with randomly chosen cluster sizes (fig. 2B). As expected, for clusters with a size of 0.001% the vast majority of the populations went extinct (fig. 2B,C). No more extinct populations were observed for clusters larger than 0.1% (fig. 2B,D). Hence, in the simulated scenario the minimum size of piRNA clusters is about 0.1%.

So far we assumed that cluster insertions have no deleterious fitness effects. It is however feasible that TE insertions in piRNA clusters also reduce host fitness. For example, cluster insertions may participate in ectopic recombination among TEs, which could lead to highly deleterious genomic rearrangements (Montgomery et al., 1991). We found that negative selection of cluster insertions only had a minor influence on the minimum fitness during an invasion and thus the extinction rate of populations (supplementary fig. S1). The minimum size of piRNA clusters was slightly larger when all TE insertions, including cluster insertions, had negative fitness effects (*x* = 0.01; supplementary fig. S1). For the reminder of the manuscript we assumed that cluster insertions have no fitness cost to the host (i.e. *x* = 0.0 for cluster insertions).

To identify regions of the parameter space where populations with small piRNA clusters are vulnerable to extinction we performed simulations with randomly chosen transposition rates (*u*) and negative effects of TEs (*x*). We followed the invasions for 5000 generations and recorded the results (fig. 3). In the absence of piRNA clusters three principal outcomes may be observed (see also (Kofler, 2019)). First populations may go extinct when *u* ≫ *x* (fig. 3A). Second populations may loose all TE insertions when negative selection against TEs is strong *x* > *u* (fig. 3A). The minimum fitness of these populations is usually close to the maximum fitness (*w*_*max*_ = 1.0) (supplementary fig. S2). Third stable TE copy numbers may be attained when the number of TEs removed by negative selection equals the number of novel TE insertions gained by transposition (transposition selection balance) (Charlesworth and Charlesworth, 1983; Charlesworth and Langley, 1989). In the absence of piRNA clusters stable TE copy numbers are solely observed in a narrow region of the parameter space (fig. 3A). If large piRNA clusters (3%) are introduced into the model extinction of populations is prevented over the entire parameter space (fig. 3B; see also Kofler (2019)). Furthermore, stable TE copy numbers are observed for a wide range of parameters (*u* > *x*; fig. 3B). However, when piRNA clusters are small (0.01%) extinct populations are reemerging (fig. 3C,D). Extinct populations are mainly observed when transposition rates are high and negative effects of TEs intermediate (fig. 3C). This raises the question why just intermediate effects of TEs lead to extinctions when piRNA clusters are small. Strongly deleterious TEs are usually quickly removed by negative selection. These TEs are thus unable to accumulate to copy numbers high enough for driving populations extinct. Weakly deleterious TEs may accumulate to high copy numbers before cluster insertions stop an invasion. However the cumulative deleterious effect of TEs with weak effects may be insufficient to drive populations extinct. Solely TEs of intermediate effects may accumulate to high numbers with effects sufficiently deleterious for driving populations extinct.

**Figure 3:**
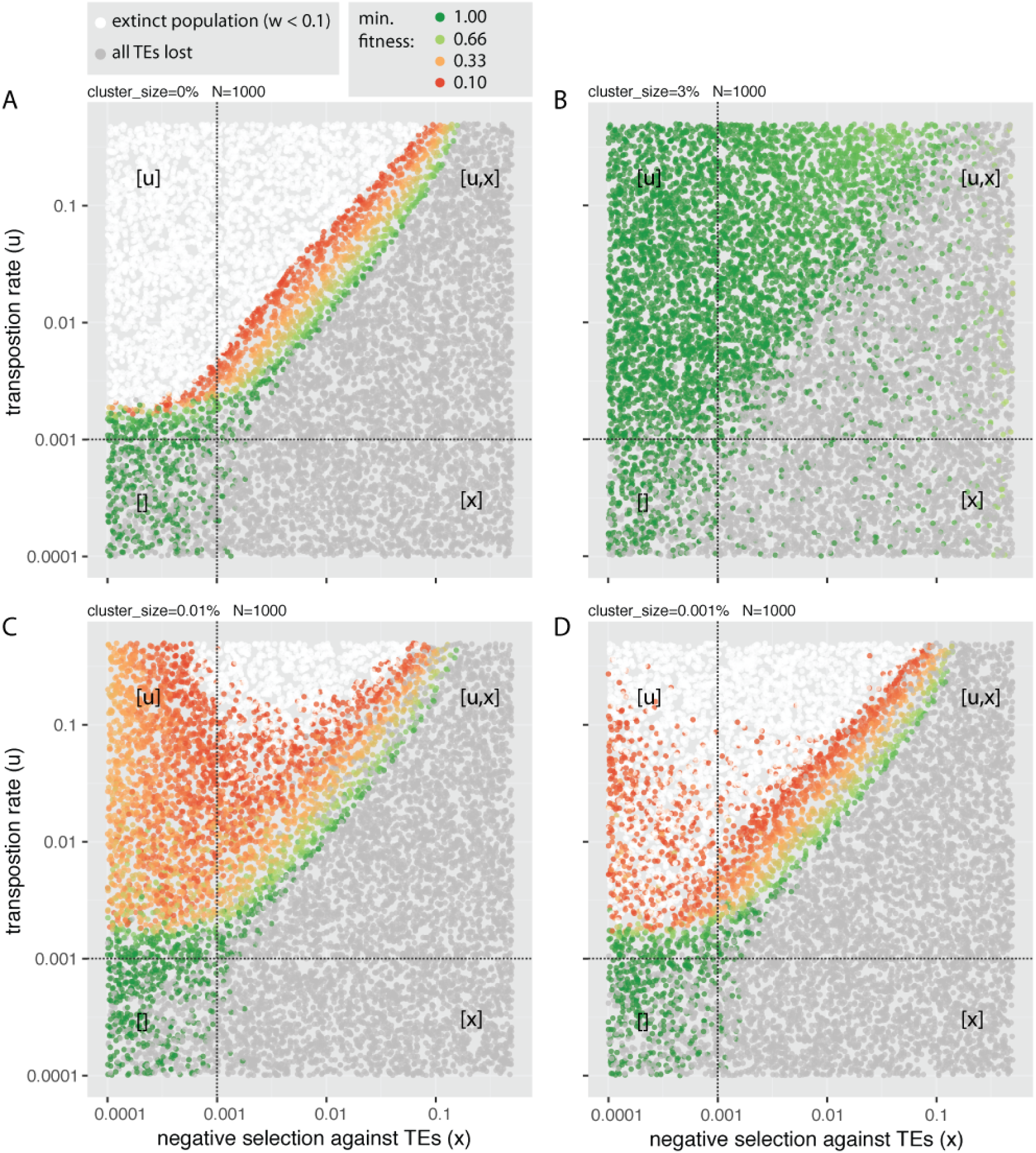
Small piRNA clusters offer an insufficient protection from extinction of host populations. Each dot represents the outcome of a single simulated TE invasion. The transposition rate (*u*) and negative effect of TEs (*x*) were randomly picked. A) In the absence of piRNA clusters extinct populations are observed when *u* ≫ *x*; B) Extinct populations are not observed when piRNA clusters are large (3% of genome). C) For small piRNA clusters (0.01%) extinct populations are observed when transposition rates are high and negative effects of TEs intermediate. D) Extinct populations are common for very small piRNA clusters (0.001%). Depending on the efficacy of negative selection and transposition (*N* * *u* > 1 and *N* * *x* > 1 with *N* = 1000) the parameter space can be divided into four quadrants. Factors that are effective in a quadrant are shown in brackets.

Next we aimed to identify the minimum size of piRNA clusters under different scenarios. We first investigated the influence of the transposition rate (*u*) and the negative effect of TEs (*x*) using >20.000 simulations with randomly chosen parameters (fig. 4). Since TE insertions are usually rapidly purged from populations when *x* > *u* (fig. 3) we performed simulations mostly with transposition rates larger than the negative effects of TEs (fig. 4). We identified regions of the parameter space that do not require piRNA clusters for controlling TEs using simulations performed without piRNA clusters (fig. 4A,B; non-white dots in left panels). piRNA clusters had a minimum size whenever piRNA clusters were necessary to control TE invasions (fig. 4A,B). The largest piRNA clusters were necessary when negative effects of TEs were intermediate (fig. 4A) and transposition rates were high (fig. 4B). The largest piRNA cluster of an extinct population had a size of 0.16%. This extinction was however observed for a simulation with a high transposition rate of *u* = 0.96. So far the largest observed transposition rates were slightly smaller, with *u* = 0.2 − 0.6 (Emilie et al., 2016; Kofler et al., 2018).

**Figure 4:**
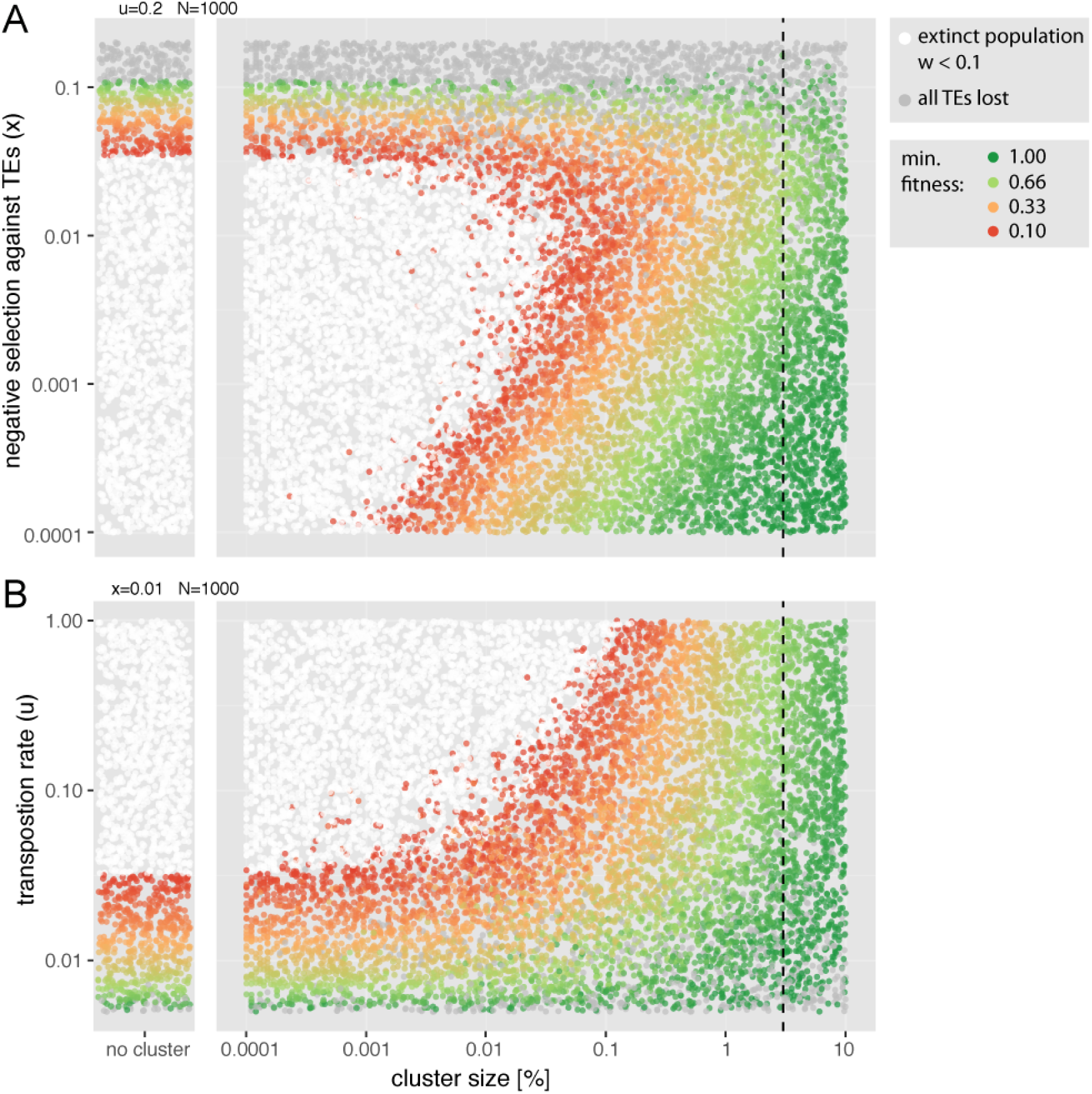
Influence of the negative effect of TEs (A) and transposition rate (B) on the minimum size of piRNA clusters. Each dot represents the outcome of a single simulation. The cluster size, the transposition rate and the negative effect of TEs were randomly picked. To identify regions of the parameter space where piRNA clusters are not necessary to control TEs (non-white dots in left panel), we show results for simulations performed without piRNA clusters. Dashed lines indicate the size of piRNA clusters in *D. melanogaster* (3%). Large piRNA clusters with sizes of 0.1-1% are necessary to control TEs when transposition rates are high and negative effects of TEs intermediate.

Since the efficacy of negative selection, an important factor counteracting TEs, depends on the population size we investigated the influence of the populations size. We performed 2000 simulations with randomly chosen cluster sizes for three different populations sizes (20000, 2000 and 200; fig. 5A). The population size had a significant influence on the minimum size of piRNA clusters (fig. 5B), where small populations required the largest piRNA clusters (fig. 5B). With the smallest evaluated population size (*N* = 200) the largest cluster of an extinct population had a size of 0.13%

**Figure 5:**
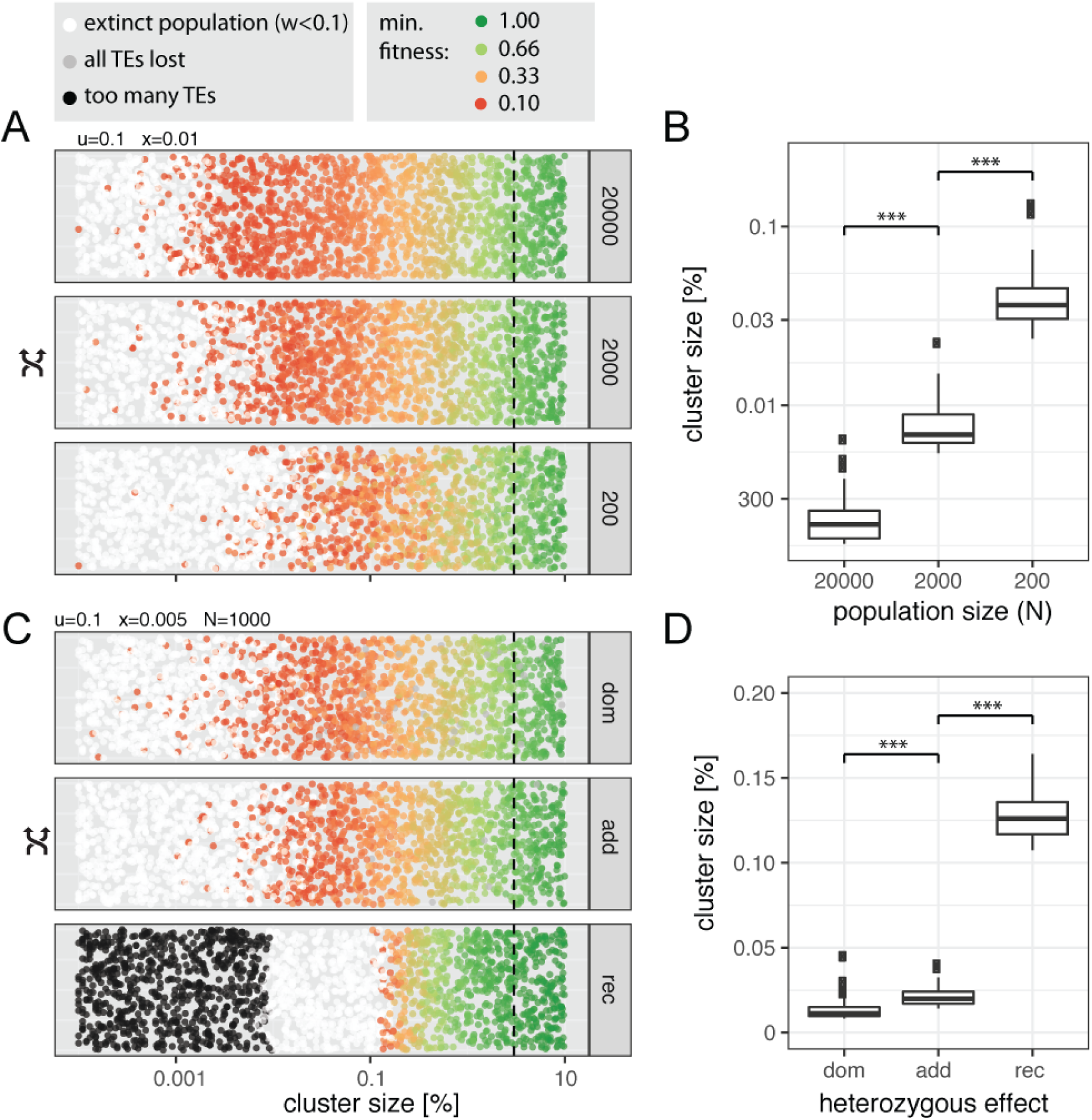
Influence of the population size (A,B) and the heterozygous effect (C,D) on the minimum size of piRNA clusters. A,C) Each dot represents the outcome of a single simulation with randomly chosen cluster sizes, population sizes (200-20000) or heterozygous effects of TEs (dom dominant, add additive, rec recessive). Dashed lines indicate the size of piRNA clusters in *D. melanogaster* (3%). B,D) The 50 largest clusters of extinct populations. Significance was estimated with Wilcoxon rank sum tests. *** *p* < 0.001.

So far we assumed that TE insertions have additive fitness effects, where homozygous TE insertions reduce host fitness twice as much as heterozygous insertions. It is however feasible that TE insertions have recessive deleterious effects. For example, TE insertions may disrupt genes for which a single intact copy is sufficient to generate the wild type (haplosufficient) (Charlesworth and Langley, 1989; Barrón et al., 2014; Lee and Langley, 2010). On the other hand it is feasible that TE insertions are dominant, for example due to trans-epigenetic effects (Lee et al., 2019). To investigate the impact of heterozygous effects on the minimum cluster size we used the fitness function *w* = 1 − 2*xi*_*hom*_ − 2*hxi*_*het*_ where *i*_*hom*_ and *i*_*het*_ are the number of homozygous and heterozygous insertion sites, respectively. For the additive case (*h* = 0.5) this equation simplifies to our standard fitness formula *w* = 1 − *xn* since the number of TE insertions per diploid genome (*n*) is *n* = 2*i*_*hom*_ + *i*_*het*_. This approach allows to model dominant (*h* = 1.0), additive (*h* = 0.5) and recessive (*h* = 0.0) effects of TE insertions. During some simulations with recessive effects, the number of accumulated TE insertions exceeded our computational resources. We terminated these simulations (> 25, 000 insertions per diploid; fig. 5C, black dots). However, since this problem solely occurred for very small piRNA clusters, an order of magnitude smaller than the clusters where extinctions were first observed, this limitation will not influence our conclusions (fig. 5C). The heterozygous effect had a significant influence on the minimum size of piRNA clusters, where the largest clusters were required for recessive insertions and the smallest for dominant insertions (fig. 5D). With recessive insertions the largest cluster of an extinct population had a size of 0.16% and with dominant insertions 0.04%. Our findings highlight the influence of the efficacy of negative selection against TEs on the minimum size of piRNA clusters. When the efficacy of negative selection is high (large populations or dominant insertions) small clusters are sufficient to control TEs whereas large clusters are necessary when the efficacy of negative selection is weak (small populations or recessive insertions).

So far we assumed that all TE insertions reduce host fitness by the same amount, irrespective of the genomic insertion site. It is however likely that different TE insertions have very diverse fitness effects. For example, insertions into coding sequences are probably more harmful than insertions into intergenic regions. We evaluated the effect of heterogenous fitness effects of TE insertions using the fitness function: 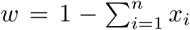. This equation again simplifies to our standard fitness formula (*w* = 1 − *xn*) when all TE insertions have identical effects. We mixed TE insertions with strong (*x* = 0.1), moderate (*x* = 0.01), weak (*x* = 0.001) and very weak (*x* = 0.0001) deleterious effects at different proportions (supplementary fig. S3A). Heterogeneity of the effect sizes had a significant influence on the minimum size of piRNA clusters (Kruskal-Wallis rank sum test with the 50 largest clusters of extinct populations; *χ*^2^ = 235, *p* < 2.2*e* − 16; supplementary fig. S3B). In all simulated scenarios piRNA clusters had a minimum size, ranging from 0.005% to 0.05% (supplementary fig. S3B).

In this work we assumed that each novel TE insertion reduces host fitness by the same amount, irrespective of the number of TE copies already present in the host genome. That is we assumed that host fitness declines linearly with TE copy numbers. Nevertheless, it is feasible that host fitness decreases more steeply with TE copy numbers. For example, ectopic recombination among distant TEs could lead to highly deleterious genomic rearrangements (e.g. inversions or translocations) and the number of ectopic recombination events may increase exponentially with TE abundance (Charlesworth and Charlesworth, 1983; Charlesworth and Langley, 1989; Montgomery et al., 1991). On the other hand, it is also conceivable that the decline of host fitness slows down with TE copy numbers. For example, the products of a few TE insertions may suffice to reduce host fitness to some extent, but the products of further TE copies may only have a minor impact. According to Charlesworth and Charlesworth (1983) we modeled interactions among TEs using the equation *w* = *xn*^*t*^. This formula allows to model a linear (*t* = 1; yielding our standard fitness formula *w* = *xn*), steep (*t* > 1) and slow (*t* < 1) decrease of host fitness with TE copy numbers (fig. 6A). We performed 10,000 simulations with randomly chosen cluster sizes and interaction terms ranging from 0.75 (slow decline) to 1.5 (steep decline; fig. 6). In the simulated scenario piRNA clusters were not necessary to control TE invasions when *t* > 1.25 (non-white dots; fig 6B). This is in agreement with previous works showing that a steep fitness decline (*t* > 1) extends the parameter space over which TEs may be controlled in the absence of piRNA clusters (Charlesworth and Charlesworth, 1983; Kofler, 2019). Nevertheless, solely large piRNA clusters protected populations from extinction over the entire parameter space (Kofler, 2019). However with less steep fitness declines (*t* < 1.25) extinct populations were observed for small piRNA clusters. Interestingly the minimum size of piRNA clusters only varied moderately among the different epistatic interactions of TEs (0.017% with *t* ≈ 0.75; 0.045% with *t* ≈ 1.0; 0.015% with *t* ≈ 1.2). Our finding that piRNA clusters have a minimum size is thus robust to different non-linear declines of host fitness with TE copy numbers.

**Figure 6:**
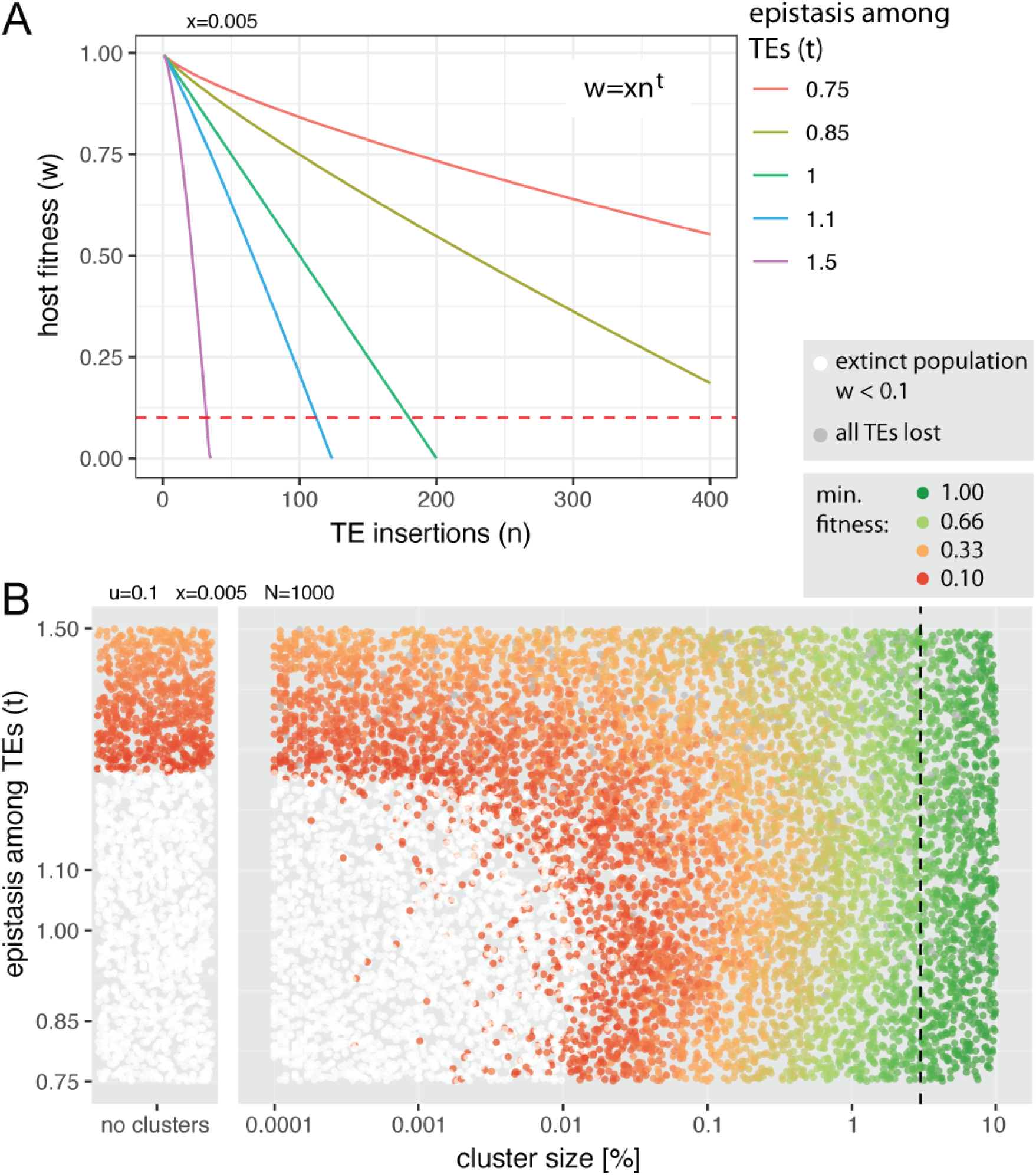
Influence of epistatic interactions (*t*) among TEs on the minimum size of piRNA clusters A) Fitness of the host may decrease linearly (*t* = 1), steeply (*t* > 1) or slowly (*t* < 1) with TE copy numbers. Red dashed line indicates the extinction threshold (*w* < 0.1). B) Parameter space with epistatic TE interactions. Each dot represents the outcome of a single simulation with a randomly picked cluster size and epistatic interaction term (*t*). To identify regions of the parameter space where piRNA clusters are not necessary to control TEs (non-white dots in left panel), we show results of simulations performed without piRNA clusters. Dashed line indicates the size of piRNA clusters in *D. melanogaster* (3%).

Under the previously simulated scenarios piRNA clusters with a size of 0.2% were sufficient to protect populations from extinction. The joint effect of some important parameters may however necessity even larger clusters. We thus investigated the minimum size of piRNA clusters under a worst case scenario, i.e: i) small population size (20-100) ii) recessive TE insertions iii) high transposition rates (*u* = 0.2) and iv) intermediate negative effects of TEs (*x* = 0.01; fig. 7). Under this scenario piRNA clusters need a size of 1-3% to protect populations from extinction (fig. 7).

**Figure 7:**
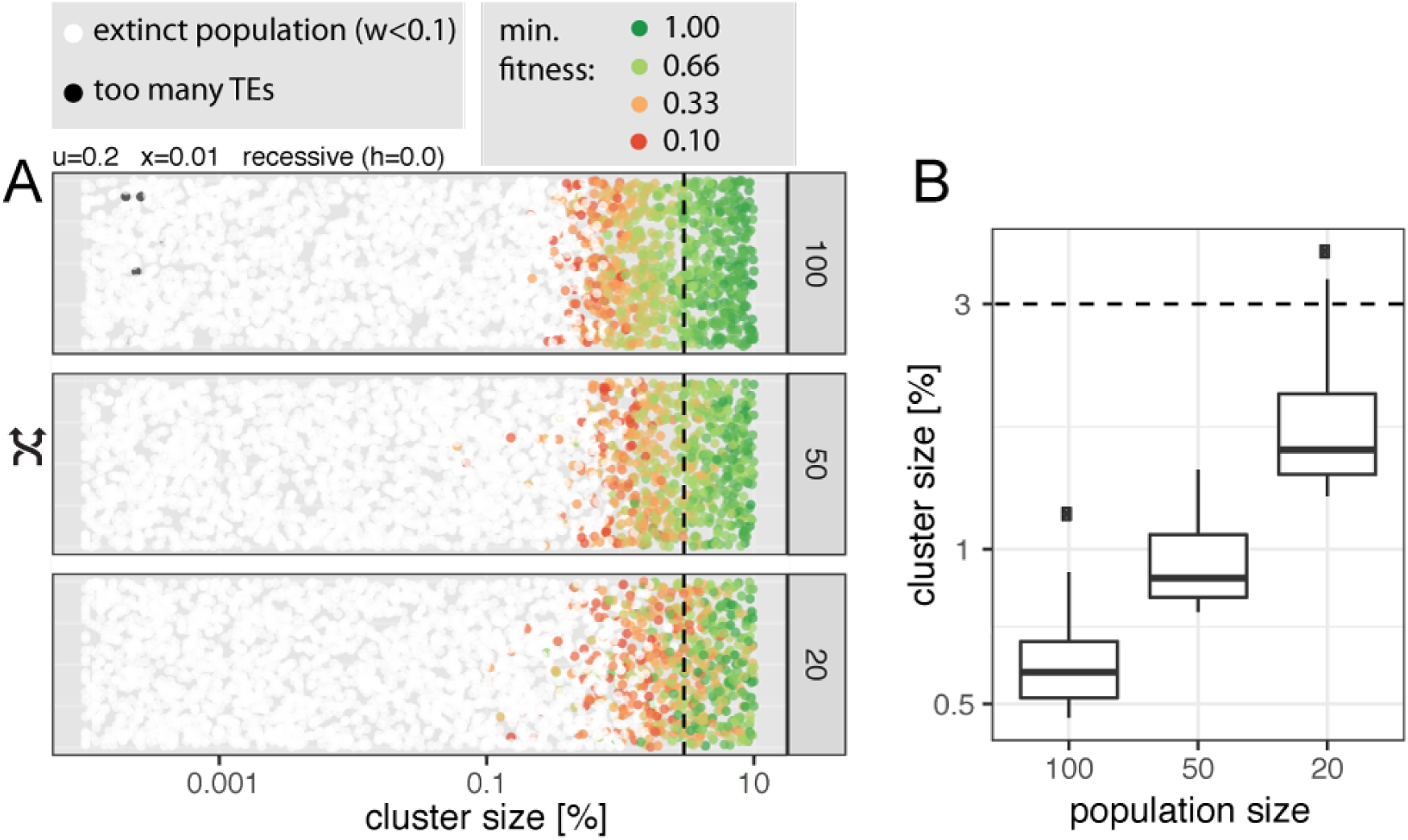
Minimum size of piRNA clusters under the worst case scenario: recessive TE insertions of intermediate effects, high transposition rates, and three different small population sizes (right panel). A) Influence of the cluster size on extinction of populations. Each dot represents the result of single simulation with a randomly drawn cluster size. The minimum fitness is shown for non-extinct populations. The size of piRNA clusters in *D. melanogaster* is shown as dashed black line (3%). B) The 50 largest piRNA clusters of extinct populations.

In summary we conclude that piRNA clusters need a size of at least 0.2% to protect populations from extinction over a wide range of parameters. Under a worst case scenario, involving small populations and recessive insertions, clusters accounting for up to 3% of genomes may be necessary to control TEs.

## Discussion

Here we showed that a simple population genetics model reveals a constraint on the size of transposons traps, such as piRNA clusters. To control TE invasions over a wide range of parameters piRNA clusters need to have a minimum size of 0.2% of a genome. When clusters are smaller populations may go extinct from the load of deleterious TE insertions accumulating during a TE invasion

Our finding that piRNA clusters have a minimum size rests on three well supported assumptions. First, TEs amplify within genomes at a certain rate. Second, since TE insertions are mostly deleterious host fitness decreases with TE copy numbers such that hosts will eventually die from the cumulative effects of the TE insertions. Third, piRNA clusters act as transposon traps, where a TE insertion in a piRNA cluster represses all other copies of the TE in trans.

There is little doubt about our assumption that TEs multiply within host genomes (Burt and Trivers, 2008; Wicker et al., 2007). However, we also relied on the widely used assumption that the transposition rate is constant during an invasion (in the absence of piRNAs) (Charlesworth and Charlesworth, 1983; Charlesworth, 1991; Kelleher et al., 2018; Lu and Clark, 2009; Kofler, 2019). It is not entirely clear if this assumption holds since very few studies estimated transposition rates during TE invasions. Kofler et al. (2018) traced P-element invasions in experimental *D. simulans* populations for 40-60 generations. Before the TE was silenced by piRNAs the P-element had a rather uniform transposition rate of about *u* = 0.15 at hot conditions and *u* = 0.05 at cold conditions (Kofler et al., 2018). Hence, a constant transposition rate is currently a reasonable assumption.

Our assumption that TE insertions are mostly deleterious is also well supported (Yukuhiro et al., 1985; Mackay, 1989; Mackay et al., 1991; Houle and Nuzhdin, 2004; Blumenstiel et al., 2014; Pasyukova et al., 2004). Moreover it seems highly unlikely that a complex host defence mechanisms, such as the piRNA pathway; could have evolved if TEs were not deleterious. Nevertheless, several major questions remain open. Most importantly, the distribution of fitness effects of TE insertions and the rate at which fitness decays with TE copy numbers remains unclear. Our finding, that piRNA clusters need a minimum size to control TE invasions, is however robust to different assumptions about the deleterious effects of TEs. Our conclusion would not hold under a model were host fitness never approaches zero, even when organism accumulate vast amounts of TEs, but such a model seems unlikely. Indeed there is ample support for our assumption that TEs may cause the death (or sterility) of hosts. The well described hybrid dysgenesis systems are good examples. Since the piRNA mediated defence against TEs is maternally transmitted, TEs are reactivated in offspring resulting from crosses of males having a TE with females not having a TE. This TE reactivation may lead to diverse defects. The I-element causes developmental defects that prevent hatching of larvae (Bucheton et al., 1976, 1992; Wang et al., 2018). Hobo and P-element activity lead to dysgenic and mostly sterile ovaries (Kidwell, 1983; Kidwell et al., 1977; Blackman et al., 1987; Hill et al., 2016). Sterile gonads may also result from reactivation of multiple TE families as for example observed with a hybrid dysgensis syndrome in *D. virilis* (Evgen’ev et al., 1997; Erwin et al., 2015). *Furthermore, individuals with defects in components of the piRNA pathway, such as Aub* or *Piwi*, are frequently infertile (Lin and Spradling, 1997; Théron et al., 2018; Czech et al., 2013). This infertility has been attributed to an upregulation of TE activity (Czech et al., 2013).

Finally our assumption that piRNA clusters act as traps controlling TE activity is well supported. First it was observed that a single *P-element* insertion in a piRNA cluster is sufficient for repressing a reporter construct in trans (Josse et al., 2007). Second, insertion of an artificial sequence into a piRNA cluster triggered production of piRNAs complimentary to the sequence (Muerdter et al., 2012). Third, the transposons ZAM and Idefix are inactive in strains having an insertion of these two TEs in the somatic piRNA cluster flamenco but active in strains not having these cluster insertions (Zanni et al., 2013). Fourth, a *de novo* insertion of ZAM into a germline piRNA cluster restores repression of ZAM (Duc et al., 2019). Fifth, piRNA cluster are mostly composed of TE sequences (Brennecke et al., 2007), supporting the view that clusters carry the trapped remnants of past TE invasions. Sixth, computer simulations confirm that genomic traps, such as piRNA clusters, may stop TE invasions (Kofler, 2019; Kelleher et al., 2018; Lu and Clark, 2009).

The minimum size of piRNA clusters inferred in this work (0.2%) rests on the further assumption that TE insertions sites are random. Some transposons however have pronounced insertion preferences (Sultana et al., 2017) that may lead to over- or underrepresentation of TE insertions within piRNA clusters. Our estimates of the minimum size of piRNA clusters will be too small if TEs avoid inserting into piRNA clusters and too large if TEs preferentially jump into piRNA clusters. Since insertion biases vary among TE families, a robust protection from extinction may require piRNA clusters larger than the estimated 0.2%.

Our estimated minimum size of piRNA clusters is solely based on a population genetics model. It is likely that biochemical processes also constrain the size of piRNA clusters. For example (de Vanssay et al., 2012) showed that loci with 7 tandem copies of a transgene may be converted into a piRNA cluster by maternally inherited small RNAs whereas loci with fewer tandem copies could not be converted.

Our finding that piRNA clusters need a minimum size of 0.2% (3% in worst case) to control TE invasions raises the important question if clusters of different species actually meet this requirement. The size of germline clusters in *D. melanogaster* is about 3% of the genome (Brennecke et al., 2007). These clusters thus provide a comprehensive protection from TEs even under a worst case scenario. Solely a TE with a strong insertion bias against piRNA clusters could overcome this defence and drive populations extinct. In *D. melanogaster* a distinct set of transposons is active in the somatic tissue surrounding the germline. These TEs are controlled by a single somatic cluster, flamenco, which has a size of about 0.15% (assuming a flamenco size of 300 kb and a genome size of 200 Mb, Brennecke personal communication Bosco et al. (2007)). As another example, in Koala piRNA cluster account for 0.17% of the genome (Yu et al., 2019). Somatic clusters in *D. melanogaster* and piRNA cluster in Koala provide sufficient protection under the majority of the simulated scenarios. However, under a worst case scenario, i.e. small populations and recessive TE insertions, these clusters may be too small to control TEs. In mice, humans and rats pachytene piRNA clusters account for about 0.1% of the genomes (Girard et al., 2006). These clusters likely offer an insufficient protection from TE invasions under multiple simulated scenarios. Nevertheless, pachytene piRNA cluster may be able to stop TE invasions if the TEs have an insertion bias into these clusters (Ernst et al., 2017). Alternatively, some organism may have multiple layers of defence against TEs. For example Kruppel-associated box zinc-finger proteins (KRAB-ZFPs) may silence TEs in some mammals (Yang et al., 2017). KRAB-ZFPs bind to TE sequences and induce heterochromatin formation (Yang et al., 2017). However, some TEs continue to be active even when targeted by KRAB-ZFPs and KRAB-ZFPs only target some TE families (Yang et al., 2017). Furthermore, it not clear if KRAB-ZFPs are mostly responsible for the long-term maintenance of TE repression or if they may also silence a newly invading TE. It is however difficult to see how the sequence specificity of KRAB-ZFPs, which is determined by tandem arrays of C2H2 zinc fingers (Yang et al., 2017), could rapidly adapt to the sequence of a newly invading TE. Thus in contrast to the piRNA pathway, which provides a generalized defence mechanism capable of trapping and silencing any mobile element, KRAB-ZFPs likely constitute a more specialized defence layer, requiring adaptation of the DNA binding sites to the sequence of each novel TE. It would be interesting to estimate the size of piRNA clusters in more species. Blatant violations of our minimum size requirements may indicate that the piRNA pathway is not the primary defence against TEs in some organism or that the simple version of the trap model, which presumes that a TE invasion is controlled by random TE insertions in piRNA clusters, is incomplete. It may for example be necessary to incorporate paramutations into the trap model (Shpiz et al., 2014; Mohn et al., 2014; Le Thomas et al., 2014; de Vanssay et al., 2012). Under this scenario maternally transmitted piRNAs may convert some euchromatic TE insertions into piRNA producing loci. Silencing of a novel TE invasion with paramutations could thus proceed like a chain reaction where increasing amounts of TE insertions in different individuals are converted into piRNA producing loci. The source of the first piRNAs that trigger this chain reaction may again be a random insertions into piRNA clusters. In contrast to the simple trap model, where several cluster insertions are required in each individual (Kofler, 2019), cluster insertions are solely expected in very few individuals under a trap model with paramutations (since euchromatic TE insertions may also yield piRNAs). The minimum size of piRNA cluster would thus be much reduced. Currently many questions about paramutations remain to be answered: i) are paramutations important for silencing novel TE invasions or mostly for maintaining silencing of a TE, ii) which fraction of the TE insertions could potentially be converted into piRNA producing loci iii) are paramutated loci stably inherited over generations and iv) what is the role of the environment in paramutations (e.g. Casier et al., 2019).

Finally, it is also feasible that in some organism piRNAs against a newly invading TE emerge independently of cluster insertions. It was for example proposed that the emergence of piRNAs against the KoRV virus in koala is triggered by unspliced TE transcripts (Yu et al., 2019). The mechanism responsible for recognizing unspliced TE transcripts is however still unclear. Furthermore it is not clear if this process may silence DNA transposons and how this mechanism may be able to distinguish between transcripts derived from TEs and genes.

Our work also raises the important question of which evolutionary forces shape the evolution of piRNA clusters. In our simulations, populations with small piRNA clusters went extinct while populations with large clusters survived. It is however not necessary to invoke such group selection arguments (Smith, 1964; Maynard Smith, 1976) to explain the evolution of piRNA clusters in natural populations. The size of piRNA clusters could be polymorphic in natural populations, such that individuals with small and large clusters can be found within a single population. If individuals with large clusters end up with fewer deleterious TE insertions than individuals with small clusters, large clusters may be positively selected. This hypothesis is supported by recent simulation studies which found that TE insertions within piRNA clusters may be positively selected (Lu and Clark, 2009; Kofler, 2019; Kelleher et al., 2018). Large clusters will on the average end up with more TE insertions than small cluster. Due to perfect linkage between piRNA clusters and its TE insertions large cluster will be positively selected. The size of piRNA clusters may thus grow to a level where extinctions of populations are no longer expected.

On the other hand evolutionary forces may exist that restrict the size of piRNA clusters. If the TE defence machinery comes at a cost to the host (Koonin et al., 2019), large piRNA clusters may be more costly to maintain than small cluster, especially when novel TE invasions are rare. The fitness cost of piRNA clusters may for example stem from ectopic recombination among cluster insertions or the cellular resources expended for generating vast amounts of piRNAs derived from large portions of genomes. This raises the intriguing possibility that the size of piRNA clusters is at an equilibrium between evolutionary forces that act to expand and contract piRNA clusters. If this hypothesis is true natural populations should exhibit variation in the size of piRNA clusters. Such variation could exist. A recent work found that the subtelmoreric piRNA cluster X-TAS, is present in most wild strains of *D. melanogaster* but frequently lost in lab strains (Asif-Laidin et al., 2017). Future studies may shed light on variation of piRNA clusters in natural populations.

## Acknowledgements

We thank Florian Schwarz for comments and all members of the Institute of Population Genetics for feedback and support. This work was supported by an Austrian Science Foundation (FWF) grant P30036-B25 to RK.

## Material and Methods

### Simulation software

All simulations were performed with the Java tool Invade (Kofler, 2019). Invade performs individual based forward simulations of TE activity in diploid organisms. For this work we released a version (0.8.07) that implements the following new features: i) compute the minimum fitness during an invasion ii) support for heterozygous effects of TE insertions and iii) support for mixed fitness effects of TE insertions. At each generation Invade performs the following steps in the given order 1) mate pairs are formed based on the fitness of the individuals, 2) haploid gametes are generated based on the recombination map, 3) novel TE insertions are introduced into the gametes of parents not having an insertion in a piRNA cluster, 4) zygotes are formed, 5) the fitness and the number of cluster insertions of the new individuals is computed and 7) the output is generated (optional).

### Simulated scenarios

We simulated a genome with five chromosomes of 10Mb (–genome Mb:10,10,10,10,10), a recombination rate of 4 cM/Mb (–rr cM Mb:4,4,4,4,4). Invasions were launched by randomly introducing 1000 TE insertions in individuals of the starting population (–basepop seg:1000). Solely simulations under our worst case scenario were launched by randomly distributing 100 insertions in the starting population. This was necessary as 1000 insertions with a *x* = 0.01 in a small population of 20 reduce the fitness of individuals in the starting population by 50% (with 100 insertions fitness is solely reduced by 5%). The transposition rate (e.g. –u 0.1), the negative effect of TE insertions (e.g. –x 0.01), the population size (e.g. –N 1000), the cluster size (e.g. –cluster kb:each:100) varied among simulated scenarios. The key parameters used for each scenario are shown at the top of the figures. All TE invasions were simulated for 5000 generations (–gen 5000), except for the scenario in which we varied the transposition rate from 0.005 to 1.0, where 10.000 generations were used. To cover the parameter space in one or two dimensions we used Python scripts that launched between 2.000 and 12.500 simulations with randomly chosen parameter combinations.

Different rates of fitness decay with increasing TE copy numbers were simulated with an epistatic interaction term interaction term (e.g. –t 1.0; with the fitness function *w* = *xn*^*t*^). To simulate dominant, recessive and additive TE effects a heterozygous effect was provided (recessive –nsmodel het:0.0; additive –nsmodel het:0.5; dominant –nsmodel het:1.0). Mixed effects of TE insertions were simulated by providing the effect sizes of the insertion sites (–nsmodel site:0.1,0.001). Equal amounts of sites where simulated for each effect size.

All statistical analysis where performed using R (R Core Team, 2012) and visualizations were done with the ggplot2 library (Wickham, 2016).

The size of piRNA clusters of humans, mouse and rat was computed by using the data of Girard et al. (2006) (supplementary tables 2-4). We summed the length of each cluster and divided this sum by the approximate genome size of 3000Mbp (cluster sizes: mouse 2.70Mbp, human 3.16Mbp, rat 3.29Mbp).

### Availability

Invade is available at https://sourceforge.net/projects/invade/. All Python and R scripts used for the analysis are available at sourceforge https://sourceforge.net/projects/te-tools/ (folder “minsize”)

## References

Arkhipova, I. R. (2018). Neutral Theory, Transposablel Elements, an Eukaryotic Genome Evolution. Mol Biol Evol, 35(6):1332–1337.

Asif-Laidin, A., Delmarre, V., Laurentie, J., Miller, W. J., Ronsseray, S., and Teysset, L. (2017). Short and long-term evolutionary dynamics of subtelomeric piRNA clusters in Drosophila. DNA Research, 25(5):459–472.

Barrón, M. G., Fiston-Lavier, A.-S., Petrov, D. A., and González, J. (2014). Population Genomics of Transposable Elements in Drosophila. Annual Review of Genetics, 48(1).

Bergman, C. M., Quesneville, H., Anxolabéhère, D., and Ashburner, M. (2006). Recurrent insertion and duplication generate networks of transposable element sequences in the *Drosophila melanogaster* genome. Genome biology, 7(11):R112.

Blackman, R. K., Grimaila, R., Macy, M., Koehler, D., and Gelbart, W. M. (1987). Mobilization of hobo elements residing within the decapentaplegic gene complex: Suggestion of a new hybrid dysgenesis system in *Drosophila melanogaster*. Cell, 49(4):497–505.

Blumenstiel, J. P. (2011). Evolutionary dynamics of transposable elements in a small RNA world. Trends in Genetics, 27(1):23–31.

Blumenstiel, J. P., Chen, X., He, M., and Bergman, C. M. (2014). An Age-of-Allele Test of Neutrality for Transposable Element Insertions. Genetics, 196(2):523–538.

Bosco, G., Campbell, P., Leiva-Neto, J. T., and Markow, T. A. (2007). Analysis of Drosophila Species Genome Size and Satellite DNA Content Reveals Significant Differences Among Strains as Well as Between Species. Genetics, 177(3):1277–1290.

Brennecke, J., Aravin, A. A., Stark, A., Dus, M., Kellis, M., Sachidanandam, R., and Hannon, G. J. (2007). Discrete small RNA-generating loci as master regulators of transposon activity in Drosophila. Cell, 128(6):1089–1103.

Brennecke, J., Malone, C. D., Aravin, A. A., Sachidanandam, R., Stark, A., and Hannon, G. J. (2008). An epigenetic role for maternally inherited piRNAs in transposon silencing. Science, 322(5906):1387–1392.

Brookfield, J. F. and Badge, R. M. (1997). Population genetics models of transposable elements. Genetica, 100(1-3):281–294.

Bucheton, A., Lavige, J., Picard, G., and L’heritier, P. (1976). Non-mendelian female sterility in *Drosophila melanogaster* : quantitative variations in the efficiency of inducer and reactive strains. Heredity, 36(3):305–314.

Bucheton, A., Vaury, C., Chaboissier, M. C., Abad, P., Pélisson, A., and Simonelig, M. (1992). I elements and the *Drosophila* genome. Genetica, 86(1-3):175–90.

Burt, A. and Trivers, R. (2008). Genes in conflict: the biology of selfish genetic elements. Belknap Press.

Casacuberta, E. and González, J. (2013). The impact of transposable elements in environmental adaptation. Molecular ecology, 22(6):1503–17.

Casier, K., Boivin, A., and Teysset, L. (2019). Epigenetic Inheritance : Implication of PIWI Interacting RNAs.

Charlesworth, B. (1991). Transposable elements in natural populations with a mixture of selected and neutral insertion sites. Genetical research, 57(2):127–34.

Charlesworth, B. and Charlesworth, D. (1983). The population dynamics of transposable elements. Genetical Research, 42(01):1–27.

Charlesworth, B. and Langley, C. H. (1989). The population genetics of *Drosophila* transposable elements. Annual review of genetics, 23:251–87.

Czech, B., Preall, J. B., McGinn, J., and Hannon, G. J. (2013). A transcriptome-wide RNAi screen in the drosophila ovary reveals factors of the germline piRNA pathway. Molecular Cell, 50(5):749–761.

Daborn, P. J., Yen, J. L., Bogwitz, M. R., Le Goff, G., Feil, E., Jeffers, S., Tijet, N., Perry, T., Heckel, D., Batterham, P., Feyereisen, R., Wilson, T. G., and Ffrench-Constant, R. H. (2002). A single P450 allele associated with insecticide resistance in Drosophila. Science, 297(5590):2253–2256.

de Vanssay, A., Bougé, A.-L., Boivin, A., Hermant, C., Teysset, L., Delmarre, V., Antoniewski, C., and Ronsseray, S. (2012). Paramutation in Drosophila linked to emergence of a piRNA-producing locus. Nature, 490(7418):112–115.

Doolittle, W. F. and Sapienza, C. (1980). Selfish genes, the phenotype paradigm and genome evolution. Nature, 284(5757):601–3.

Duc, C., Yoth, M., Jensen, S., Mouniée, N., and Bergman, C. M. (2019). Trapping a somatic endogenous retrovirus into a germline piRNA cluster immunizes the germline against further invasion. Genome Biology, 20(1):127.

Emilie, R., Rouzic, A. L., Zhang, Z., Capy, P., and Hua-Van, A. (2016). Experimental evolution reveals hyperparasitic interactions among transposable elements. PNAS, 113(53):14763–14768.

Ernst, C., Odom, D. T., and Kutter, C. (2017). The emergence of piRNAs against transposon invasion to preserve mammalian genome integrity. Nature Communications, 8(1):1–9.

Erwin, A. A., Galdos, M. A., Wickersheim, M. L., Harrison, C. C., Marr, K. D., Colicchio, J. M., and Blumenstiel, J. P. (2015). piRNAs Are Associated with Diverse Transgenerational Effects on Gene and Transposon Expression in a Hybrid Dysgenic Syndrome of D. virilis. PLoS genetics, 11(8):e1005332.

Evgen’ev, M. B., Zelentsova, H., Shostak, N., Kozitsina, M., Barskyi, V., Lankenau, D. H., and Corces, V. G. (1997). Penelope, a new family of transposable elements and its possible role in hybrid dysgenesis in *Drosophila virilis*. PNAS, 94(1):196–201.

Girard, A., Sachidanandam, R., Hannon, G. J., and Carmell, M. A. (2006). A germline-specific class of small RNAs binds mammalian Piwi proteins. Nature, 442(7099):199–202.

González, J., Lenkov, K., Lipatov, M., Macpherson, J. M., and Petrov, D. A. (2008). High rate of recent transposable element–induced adaptation in *Drosophila melanogaster*. PLoS biology, 6(10):e251.

Goriaux, C., Théron, E., Brasset, E., and Vaury, C. (2014). History of the discovery of a master locus producing piRNAs: The flamenco/COM *Drosophila melanogaster*. Frontiers in Genetics, 5:1–8.

Gunawardane, L. S., Saito, K., Nishida, K. M., Miyoshi, K., Kawamura, Y., Nagami, T., Siomi, H., and Siomi, M. C. (2007). A slicer-mediated mechanism for repeat-associated siRNA 5’ end formation in Drosophila. Science, 315(5818):1587–1590.

Hill, T., Schlötterer, C., and Betancourt, A. J. (2016). Hybrid Dysgenesis in *Drosophila simulans* Associated with a Rapid Invasion of the P-Element. PLoS Genet, 12(3):e1005920.

Houle, D. and Nuzhdin, S. V. (2004). Mutation accumulation and the effect of copia insertions in *Drosophila melanogaster*. Genetics Research, 83(1):7–18.

Josse, T., Teysset, L., Todeschini, A.-L., Sidor, C. M., Anxolabéhère, D., and Ronsseray, S. (2007). Telomeric trans-silencing: an epigenetic repression combining RNA silencing and heterochromatin formation. PLoS Genetics, 3(9):1633–43.

Kelleher, E. S., Azevedo, R. B. R., and Zheng, Y. (2018). The Evolution of Small-RNA-Mediated Silencing of an Invading Transposable Element. Genome Biology and Evolution, 10(11):3038–3057.

Kidwell, M. G. (1983). Evolution of hybrid dysgenesis determinants in *Drosophila melanogaster*. Proceedings of the National Academy of Sciences, 80(6):1655–1659.

Kidwell, M. G., Kidwell, J. F., and Sved, J. A. (1977). Hybrid dysgenesis in *Drosophila melanogaster* : A syndrome of aberrant traits including mutations, sterility and male recombination. Genetics, 86(4):813–833.

Kofler, R. (2019). Dynamics of transposable element invasions with piRNA clusters. Molecular Biology and Evolution, 36(7):1457—-1472.

Kofler, R., Senti, K.-A., Nolte, V., Tobler, R., and Schlötterer, C. (2018). Molecular dissection of a natural transposable element invasion. Genome Research, 28(2):824–835.

Koonin, E. V., Makarova, K. S., Wolf, Y. I., and Krupovic, M. (2019). Evolutionary entanglement of mobile genetic elements and host defence systems: guns for hire. Nature Reviews Genetics.

Le Rouzic, A. and Capy, P. (2005). The first steps of transposable elements invasion: Parasitic strategy vs. genetic drift. Genetics, 169(2):1033–1043.

Le Thomas, A., Rogers, A. K., Webster, A., Marinov, G. K., Liao, S. E., Perkins, E. M., Hur, J. K., Aravin, A. A., and Tóth, K. F. (2013). Piwi induces piRNA-guided transcriptional silencing and establishment of a repressive chromatin state. Genes and Development, 27(4):390–399.

Le Thomas, A., Stuwe, E., Li, S., Du, J., Marinov, G., Rozhkov, N., Chen, Y. C. A., Luo, Y., Sachidanandam, R., Toth, K. F., Patel, D., and Aravin, A. A. (2014). Transgenerationally inherited piRNAs trigger piRNA biogenesis by changing the chromatin of piRNA clusters and inducing precursor processing. Genes and Development, 28(15):1667–1680.

Lee, Y. C. G. and Langley, C. H. (2010). Transposable elements in natural populations of *Drosophila melanogaster*. Philosophical transactions of the Royal Society of London. Series B, Biological sciences, 365(1544):1219–28.

Lee, Y. C. G., Ogiyama, Y., Martins, N. M., Beliveau, B. J., Acevedo, D., Cavalli, G., Karpen, G. H., et al. (2019). Pericentromeric heterochromatin is hierarchically organized and spatially contacts h3k9me2/3 islands located in euchromatic genome. bioRxiv, page 525873.

Lin, H. and Spradling, A. (1997). A novel group of pumilio mutations affects the asymmetric division of germline stem cells in the Drosophila ovary. Development (Cambridge, England), 124(12):2463–2476.

Lu, J. and Clark, A. G. (2009). Population dynamics of PIWI-RNAs (piRNAs) and their targets in Drosophila. Genome Research, 20:212–227.

Mackay, T. F. (1989). Transposable elements and fitness in *Drosophila melanogaster*. Genome, 31(1):284–295.

Mackay, T. F., Lyman, R. F., and Jackson, M. S. (1991). Effects of P Element Insertions on Quantitative Traits in *Drosophila melanogaster*. Genetics, 130:315–332.

Malone, C. D. and Hannon, G. J. (2009). Small RNAs as Guardians of the Genome. Cell, 136(4):656–668.

Maynard Smith, J. (1976). Group Selection. The quarterly Review of Biology, 51(June):277–283.

Mohn, F., Sienski, G., Handler, D., and Brennecke, J. (2014). The rhino-deadlock-cutoff complex licenses noncanonical transcription of dual-strand piRNA clusters in Drosophila. Cell, 157(6):1364–1379.

Montgomery, E. A., Huang, S. M., Langley, C. H., and Judd, B. H. (1991). Chromosome rearrangement by ectopic recombination in *Drosophila melanogaster* : genome structure and evolution. Genetics, 129(4):1085–98.

Muerdter, F., Olovnikov, I., Molaro, A., Rozhkov, N. V., Czech, B., Gordon, A., Hannon, G. J., and Aravin, A. A. (2012). Production of artificial piRNAs in flies and mice. Rna, 18(1):42–52.

Nuzhdin, S. V. (1999). Sure facts, speculations, and open questions about the evolution of transposable element copy number. Genetica, 107(1-3):129–137.

Orgel, L. E. and Crick, F. H. (1980). Selfish DNA: the ultimate parasite. Nature, 284(5757):604–7.

Ozata, D. M., Gainetdinov, I., Zoch, A., O’Carroll, D., and Zamore, P. D. (2018). PIWI-interacting RNAs: small RNAs with big functions. Nature Reviews Genetics, 20(2):89–108.

Pasyukova, E., Nuzhdin, S., Morozova, T., and Mackay, T. (2004). Accumulation of transposable elements in the genome of *Drosophila melanogaster* is associated with a decrease in fitness. Journal of Heredity, 95(4):284–290.

R Core Team (2012). R: A Language and Environment for Statistical Computing. R Foundation for Statistical Computing, Vienna, Austria. ISBN 3-900051-07-0.

Shpiz, S., Ryazansky, S., Olovnikov, I., Abramov, Y., and Kalmykova, A. (2014). Euchromatic transposon insertions trigger production of novel pi-and endo-siRNAs at the target sites in the Drosophila germline. PLoS Genet, 10(2):e1004138.

Sienski, G., Dönertas, D., and Brennecke, J. (2012). Transcriptional silencing of transposons by Piwi and maelstrom and its impact on chromatin state and gene expression. Cell, 151(5):964–980.

Smith, J. M. (1964). Group Selection and Kin Selection. Nature, 201(4924):1145–1147.

Sultana, T., Zamborlini, A., Cristofari, G., and Lesage, P. (2017). Integration site selection by retroviruses and transposable elements in eukaryotes. Nature Reviews Genetics.

Théron, E., Maupetit-Mehouas, S., Pouchin, P., Baudet, L., Brasset, E., and Vaury, C. (2018). The interplay between the argonaute proteins piwi and aub within Drosophila germarium is critical for oogenesis, piRNA biogenesis and TE silencing. Nucleic Acids Research, 46(19):10052–10065.

Wang, L., Dou, K., Moon, S., Tan, F. J., Zhang, Z. Z. Z., Wang, L., Dou, K., Moon, S., Tan, F. J., and Zhang, Z. Z. Z. (2018). Hijacking Oogenesis Enables Massive Propagation Article Hijacking Oogenesis Enables Massive Propagation of LINE and Retroviral Transposons. Cell, 174(5):1082–1094.

Wicker, T., Sabot, F., Hua-Van, A., Bennetzen, J. L., Capy, P., Chalhoub, B., Flavell, A., Leroy, P., Morgante, M., Panaud, O., et al. (2007). A unified classification system for eukaryotic transposable elements. Nature Reviews Genetics, 8(12):973–982.

Wickham, H. (2016). ggplot2: Elegant Graphics for Data Analysis. Springer-Verlag New York.

Yamanaka, S., Siomi, M. C., and Siomi, H. (2014). piRNA clusters and open chromatin structure. Mobile DNA, 5(1):22.

Yang, P., Wang, Y., and Macfarlan, T. S. (2017). The Role of KRAB-ZFPs in Transposable Element Repression and Mammalian Evolution. Trends in Genetics, 33(11):871–881.

Yu, T., Koppetsch, B., Chappell, K., Pagliarani, S., Johnston, S., Silverstein, N. J., Luban, J., Weng, Z., and Thauerkauf, W. E. (2019). The piRNA Response to Retroviral Invasion of the Koala Genome. Cell, pages 1–12.

Yukuhiro, B. Y. K., Harada, K., and Mukai, T. (1985). Viability mutations induced by the P elements in *Drosophila melanogaster*. Jpn. J. Genet., 60:531–537.

Zanni, V., Eymery, A., Coiffet, M., Zytnicki, M., and Luyten, I. (2013). Distribution, evolution, and diversity of retrotransposons at the flamenco locus reflect the regulatory properties of piRNA clusters. PNAS, 110(49):19842–19847.

