## supplement for "piRNA clusters need a minimum size to control transposable element invasions"

November 11, 2019

### Supplementary figures

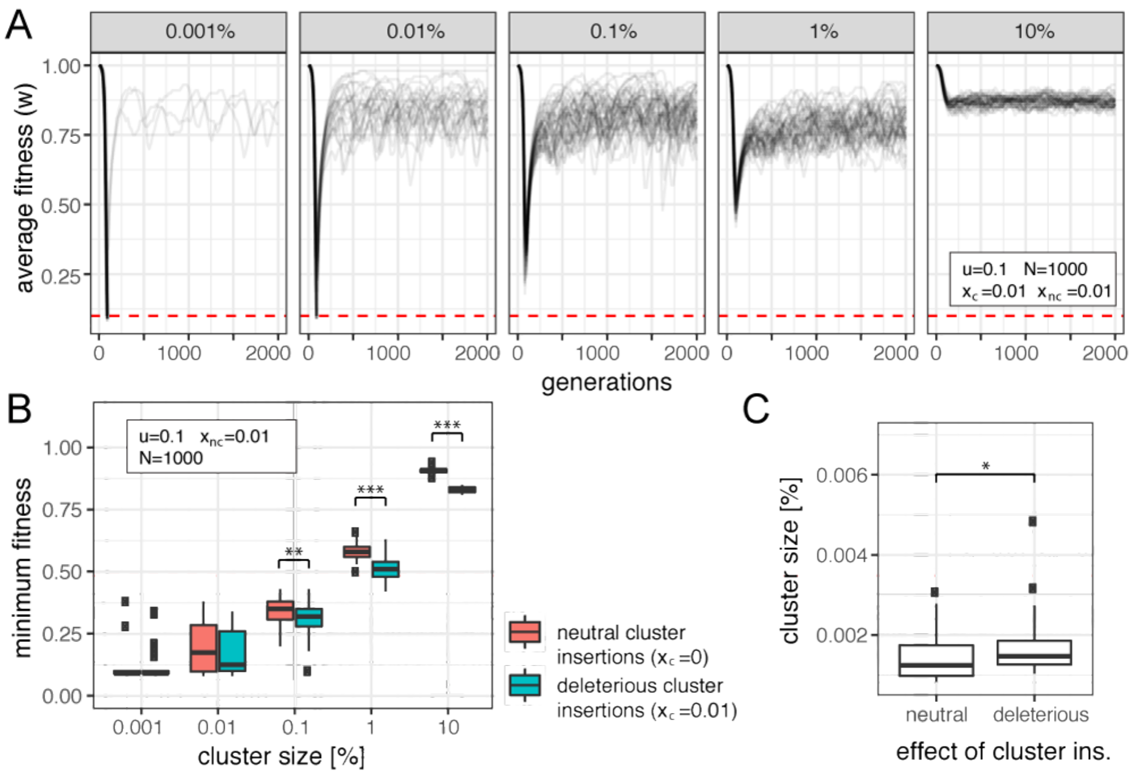


Figure 1: Impact of negative selection against cluster insertions on the minimum size of piRNA clusters. A) Average fitness of populations during TE invasions when both cluster and non-cluster insertions are negatively selected ($x_c = x_{nc}$, where $x_c$ and $x_{nc}$ is the negative effect of cluster and non-cluster insertions, respectively). Results are shown for different cluster sizes (top panel). After an initial fitness reduction, populations enter transposition-selection-cluster (TSC) balance, an equilibrium state where both negative selection and cluster insertions counteract the spread of a TE [see Kofler (2019)]. B) Effect of negative selection against cluster insertions on the minimum fitness during an invasion. C) Effect of negative selection against cluster insertions on the minimum size of piRNA clusters. Boxplots show the 50 largest clusters of extinct populations. 2000 simulations were performed for each scenario. Note that slightly larger clusters were required when cluster insertions are negatively selected. Significance was calculated with Wilcoxon rank sum tests *** $p < 0.001$, ** $p < 0.01$, * $p < 0.05$

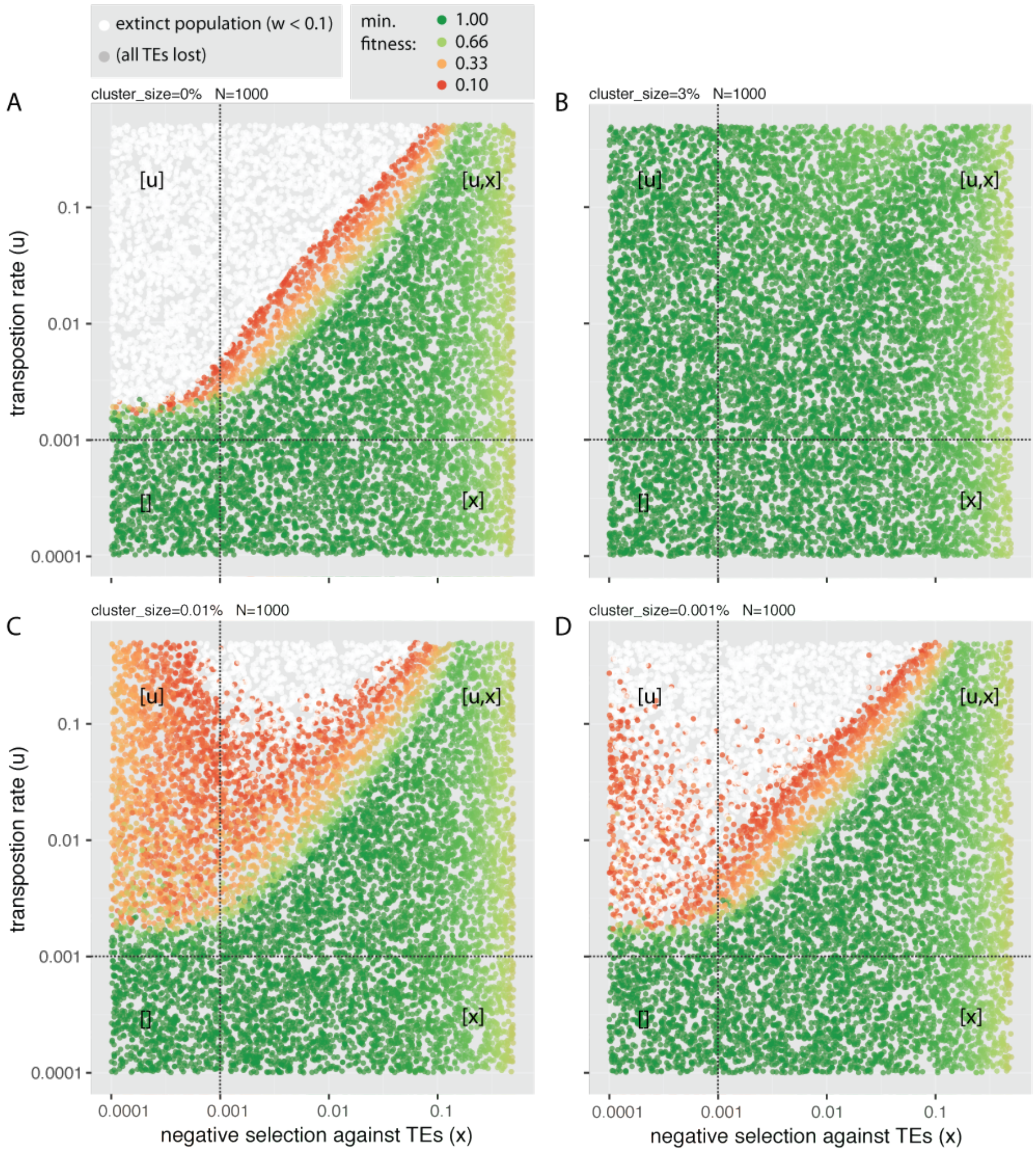


Figure 2: Parameter space showing the minimum fitness of populations for different piRNA cluster sizes (A,B,C,D). Extinct populations ($w < 0.1$) are shown separately. The minimum fitness is also shown for populations that lost all TE insertions. Each dot represents the outcome of a single simulated TE invasion. The transposition rate ($u$) and negative effect of TEs ($x$) were randomly picked. Depending on the efficacy of negative selection and transposition ($N * u > 1$ and $N * x > 1$ with $N = 1000$) the parameter space can be divided into four quadrants. Factors that are effective in a quadrant are shown in brackets.

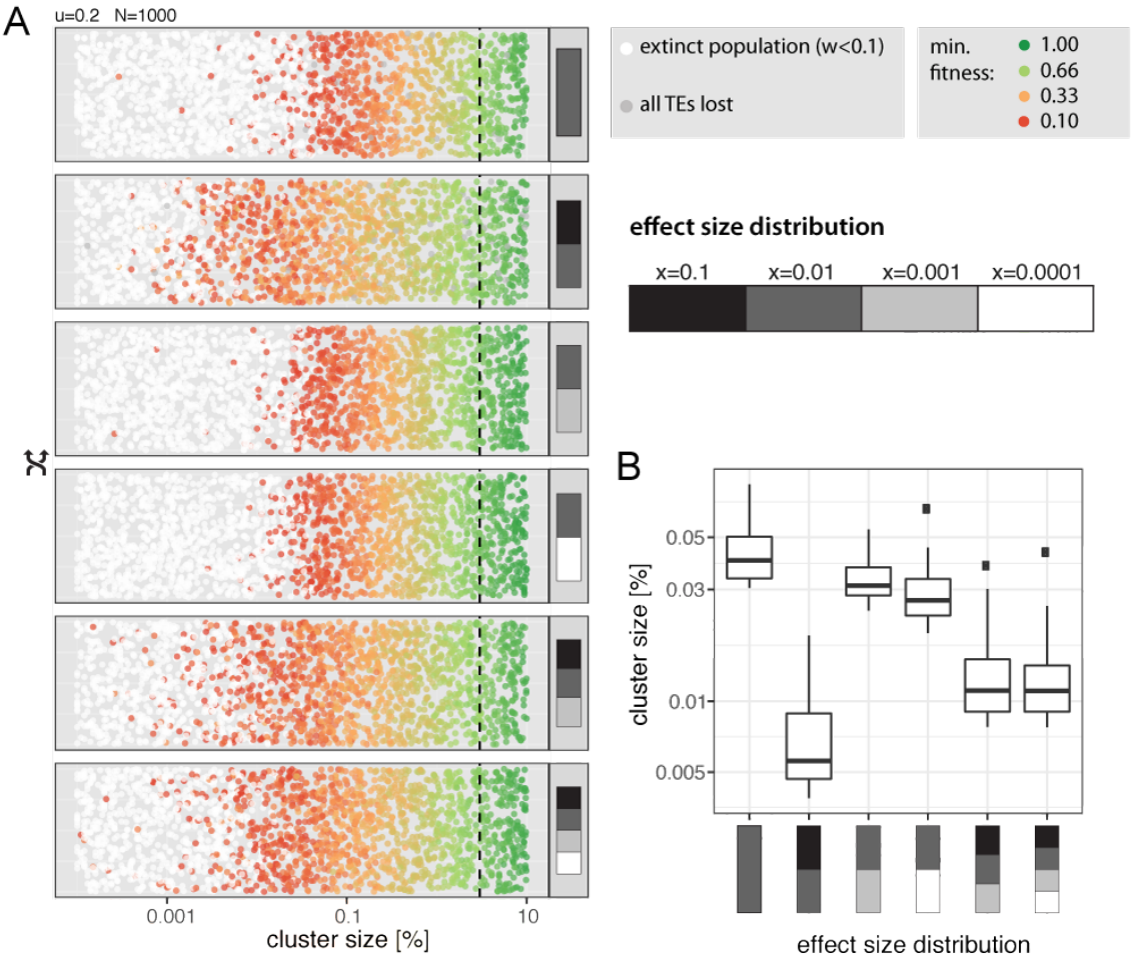


Figure 3: Influence of mixing TEs with different deleterious effects on the minimum size of piRNA clusters A) Host fitness with different mixes of TE effects (right panel). Each dot represents the result of single simulation with a randomly drawn cluster size. The minimum fitness is shown for non-extinct populations. The size of piRNA clusters in *D. melanogaster* is shown as dashed black line (3%). Mixes of TE effects are described by bars (from black and white) summing to a frequency of 1.0. Each block represents the fraction of TE insertions having the given effect size. B) The 50 largest piRNA clusters of extinct populations for different mixes of TE effects.

### References

Kofler, R. (2019). Dynamics of transposable element invasions with piRNA clusters. *Molecular Biology and Evolution*, 36(7):1457—-1472.
